# Evaluation of the Translational Readiness of Pluripotent Stem Cell–Derived Vascular Cell Therapy for Limb Ischemia: A Systematic Review and Meta-analysis

**DOI:** 10.64898/2026.09.02.749027

**Authors:** Minwoo Shin, Hyunsik Choi, Taewi Kim, Jin Pyeong Jeon, Sanghoon Jung, Kyuwon Cho

## Abstract

**Background:** For the first time, pluripotent stem cell (PSC)-derived endothelial cells were administered to a human for peripheral artery disease (PAD), in August 2026. This translation is based on a large preclinical literature claiming that donor cells build blood vessels in animal models of limb ischemia, but whether it supports that claim has not been assessed systematically.

**Methods:** In this systematic review and meta-analysis, we searched MEDLINE, Web of Science, Europe PMC, and Embase to August 2026 for studies of PSC-derived vascular cells in animal models of limb ischemia with an acellular control. Limb perfusion at each experiment’s primary timepoint was pooled as Hedges’ g in a three-level random-effects model with cluster-robust variance. Donor-cell fate and a seven-level ordinal of vascular contribution were graded for every study. This meta-analysis was registered on PROSPERO (CRD420261465308).

**Results:** Sixty-eight studies in 69 reports met the criteria; 51 contributed 114 comparisons at primary timepoints. Cell therapy increased perfusion (Hedges’ g 2.26, 95% CI 1.72–2.79; P=4.1×10-11; I2=87.5%; 95% prediction interval −1.47 to 5.98). No prespecified moderator survived Holm–Bonferroni adjustment, including the donor-cell fate grade (P=0.30). A donor cell was resolved in a vessel wall in 35 studies, and flow through such a vessel in 7. Small-study effects were strong (Egger t=6.25, P=9×10-8) but were not reproduced by a sample-size–based test (P=0.80); trim-and-fill moved the estimate to 1.15. On the natural scale, treated limbs recovered about twice the control perfusion (ratio of means 1.96, 95% CI 1.66– 2.33). Limb preservation favored cell therapy in 11 studies with countable events (risk ratio 4.88, 95% CI 2.28–10.45), but 43 of 68 studies reported no functional outcome, and only 22 comparisons from 8 studies were both randomized and blindly assessed. In a post hoc analysis that did not survive multiplicity adjustment, effects were largest in studies whose images could not resolve their incorporation claims (g 3.23) versus those that resolved incorporation (g 2.33).

**Conclusions:** PSC-derived vascular cell therapy improved limb perfusion in animal models, consistently across every sensitivity analysis, although the magnitude is uncertain given heterogeneity and small-study effects. Benefit was not associated with the degree of donor-cell incorporation across studies and was largest where incorporation was least resolvable, a pattern more consistent with reporting and measurement bias than with a true mechanistic effect. The first rigorous, blinded test of that benefit may come not from animals but from the patients now receiving these cells.

**Clinical Perspective:** *What Is New?:*

- This is the first meta-analysis of preclinical pluripotent stem cell (PSC)-derived vascular cell therapy for limb ischemia, completed as the first patients begin receiving these cells.
- Across 68 studies, PSC-derived vascular cells consistently improved limb perfusion (Hedges’ g 2.26), but the true magnitude is uncertain because the apparent small-study effect is partly a measurement artifact.
- The magnitude of benefit was not associated with the degree of donor-cell incorporation across studies (P=0.30) and was largest where structural incorporation was least resolvable, a pattern more consistent with reporting and measurement bias than with a true vascular-building mechanism.
- Donor cells were resolved within a vessel wall in 35 of 68 studies but blood flow through such a vessel in only 7, so engraftment is often shown while participation in a perfused conduit rarely is.

*What Are the Clinical Implications?:*

- Clinical translation has already begun, so effect sizes taken from this literature should not be used to power a trial, because neither the pooled estimate nor its trim-and-fill adjustment provides a reliable planning value.
- Because benefit did not depend on structural incorporation, potency assays and release criteria can be based on functional perfusion rather than on vascular incorporation alone.
- Blinded outcome assessment, demonstration of flow through donor-containing vessels in the ischemic limb, and testing in aged or comorbid animals would make future preclinical studies directly informative; yet the decisive, blinded test of benefit may now come from the patients already receiving these cells.

## Introduction

Chronic limb-threatening ischemia (CLTI) is the end stage of peripheral artery disease (PAD) and carries a high risk of major amputation and death that persists after revascularization.^1^ Median survival after a diagnosis of chronic limb-threatening ischemia is approximately 3.5 years, and is shorter still in patients whose first treatment is major amputation.^2^ Revascularization is not technically feasible or not durable in a substantial proportion of patients, and cell therapy has been pursued for more than 2 decades as an alternative for these patients. However, randomized, placebo-controlled trials of autologous bone marrow-derived and related cell products have not shown convincing benefit in major amputation, survival, or amputation-free survival, although improvement in selected surrogate measures has been reported.^3, 4^ The Global Vascular Guidelines accordingly recommend that regenerative therapies for CLTI be investigated only within rigorous clinical trials.^1^

These failures have been attributed to the cell source rather than to the concept of therapeutic angiogenesis. Pluripotent stem cells (PSCs) provide a scalable, quality-controlled, and genetically defined source of vascular cells, and their quality does not depend on the age, comorbidities, or vascular dysfunction of the patient. PSC-derived endothelial cells, vascular progenitors, pericytes, and vascular organoids have been tested in a widening range of animal models of limb ischemia, and clinical translation has now begun. In August 2026, a patient with peripheral artery disease received autologous induced pluripotent stem cell (iPSC)-derived endothelial cells under Korea’s advanced regenerative medicine clinical research framework.^5^ This is the first reported administration of a PSC-derived endothelial cell product in humans. The preclinical literature that supports this clinical translation is uniformly positive.

This uniformity warrants examination for two reasons. First, animal studies are susceptible to publication bias and to other sources of effect size inflation, and such biases account for approximately a third of the reported efficacy in experimental stroke.^6^ However, whether the preclinical literature on PSC-derived vascular cell therapy is similarly distorted has not been assessed. Second, the rationale for using vascular cells rather than any other paracrine-competent cell is mechanistic. Their vascular identity is presumed to allow donor cells to participate directly in the formation and repair of vessels, and this premise generates a testable prediction. If structural incorporation of donor cells into the host vasculature contributes to efficacy, studies demonstrating convincing incorporation should report greater functional benefit than studies in which incorporation is absent or unresolved. If the benefit is comparable regardless of incorporation, the advantage of a vascular product lies in paracrine support rather than in reconstruction of the vascular compartment. However, evidence that donor cells express endothelial markers or form vessel-like structures is not evidence that they form the wall of a perfused, host-connected vessel in the ischemic limb, and this distinction has not been evaluated systematically across the literature.

In this study, we conducted a systematic review and meta-analysis of preclinical studies of PSC-derived vascular cell therapy in animal models of limb ischemia, and we did two things that a conventional efficacy meta-analysis does not. First, we pooled the perfusion data and tested the robustness of the pooled estimate against small-study effects and other sources of bias. Second, we extracted and adjudicated what each study demonstrated about the fate of the donor cells, recording the detection method, whether donor cells were quantified, whether a donor cell was resolved within a vessel wall, whether that wall also contained host cells, and whether the evidence was obtained in the ischemic limb or in a plug assay in a healthy animal. We then asked whether the mechanistic evidence predicts the functional effect. Overall, this study evaluates the efficacy of PSC-derived vascular cell therapy in preclinical models of limb ischemia and, separately, whether the mechanism claimed to produce that efficacy is supported by the literature that claims it.

## Methods

### Transparency and data availability

The protocol was registered on PROSPERO (CRD420261465308). The complete dataset, extraction forms, risk-of-bias assessments, and analysis codes are available at https://doi.org/10.17605/OSF.IO/KG74E, in accordance with the AHA journals’ implementation of the Transparency and Openness Promotion guidelines. As a secondary analysis of already-published animal studies, the review involved no new animal experiments and did not require institutional animal-ethics approval. Amendments to the registered protocol, and deviations from it, are recorded in the protocol deviation log deposited at OSF.

### Review procedures

Two authors independently screened records on title and abstract, assessed full texts for eligibility, extracted data, digitized published figures, assessed risk of bias and reporting quality, and graded donor-cell fate. Disagreements were resolved by consensus.

### Eligibility criteria

We included controlled studies in any animal species in which a pluripotent stem cell-derived vascular cell product (endothelial, mural, pericytic, vascular-progenitor, or vascular-organoid) was transplanted in a model of hind-limb or forelimb ischemia, alongside an acellular control arm, with a quantitative measure of limb perfusion. Pluripotent sources comprised embryonic stem cells, induced pluripotent stem cells, and somatic-cell nuclear transfer-derived cells, of any species. We excluded studies without limb ischemia, without an acellular comparator, without a perfusion outcome, and reports describing animals already reported elsewhere (see *Unit of analysis* section below). Mesenchymal stromal cell (MSC) products, including MSC-like cells derived from pluripotent sources, were excluded as non-vascular lineages^7–10^. Undifferentiated PSCs were eligible only as comparators, not as index interventions. These criteria reflect the final registered protocol; the amendment history and the results under the original criteria, retained as a sensitivity analysis, are deposited at OSF. Borderline products were retained when the delivered population was defined by a vascular or vasculogenic phenotype rather than a stromal one. Sorted SSEA-1^+^/MesP1^+^ and FLK-1^+^ mesodermal vascular progenitors, pericyte-like mural cells, and the vasculogenic mesoderm product VSC100 fall within the registered vascular-progenitor and mural categories. Two studies co-delivering an adult-derived supporting population alongside the PSC-derived vascular product were retained because the PSC-derived vascular cells are the index intervention and each study provides an acellular control for that comparison.

### Search

MEDLINE via PubMed, Web of Science Core Collection, and Europe PMC were searched from inception to August 2026. Embase was searched via Ovid over the same period. Full search strategies are in the repository and are reported in line with the PRISMA-S extension for search reporting.^11^

### Study selection and data extraction

Records were screened on title and abstract against the registered criteria, and full texts of all potentially eligible records were assessed. Extraction used a predefined form covering animal, model, product, dose, route, vehicle, comparator, timepoints, sample sizes, and outcome values. Outcome values were taken from the text where reported numerically (19 of 68 studies, 5 of them only in part) and digitized from published figures where they were not (45 studies, including those 5), with each digitized series recording the group size and the error-bar type read from the legend. The remaining 9 studies yielded no usable perfusion value: 4 report no perfusion outcome at all, 4 report perfusion only in a figure that could not be read quantitatively, and 1 shows it only as a dot plot without a usable mean and dispersion.

Every perfusion series was digitized a second time, independently and blind to the first reading, and agreement was near-complete (645 matched values from the two readings; median relative difference 0.0%; Lin concordance 0.999 on the ratio scale and 0.98 on the percent scale). Values added by a later completeness check were read twice by procedures that differ rather than twice by the same procedure, once visually against the printed axis and once by an automated reading in which the axis calibration and the marker coordinates were computed from pixel positions; across 46 matched points from seven figure panels the median relative difference was 0.8% and the largest 10.0%, the latter on a value of about 2 on a 0-60 scale, and against the plotted axis range the median difference was 0.4%. Discrepant series were adjudicated to the mean of the two readings.

### Unit of analysis

Following PRISMA 2020^12^, the unit of counting is the study rather than the report. One pair of reports describes a single cohort and is counted once.^13, 14^ Six papers run the same cells in more than one host or disease model. These are recorded as separate experiments within the study, and an intervention arm is paired only with the control of its own experiment, while clustering for the multilevel model remains at the study level. Where an intervention arm receives a co-administered agent, the comparator is the acellular arm containing that agent rather than plain vehicle, which is the conservative choice.

### Risk of bias and reporting quality

Risk of bias was assessed with SYRCLE’s tool for animal studies across its ten domains.^15^ Reporting quality used a ten-item checklist adapted from CAMARADES.^16^

### Donor-cell fate and vascular contribution

A five-level donor-cell fate grade (0–4) was prespecified. As a protocol addition, we extracted five further items for every study: whether donor-cell tracking was attempted; the detection method against a controlled vocabulary; whether donor cells were quantified; the vascular contribution of donor cells on a seven-level ordinal (not assessed < donor cells not in vessels < perivascular only < incorporation claimed but not resolvable < mosaic with host cells < independent donor-only vessels < both); and whether a biomaterial carried the cells in the ischemia experiment, kept distinct from a subcutaneous plug assay in a healthy animal.

The ordinal is used for description only; for the post hoc moderator its levels are collapsed into three unordered strata (no incorporation shown, claimed but not resolvable, and resolved incorporation).

Three rules governed grading: vascular contribution was graded from the ischemic-limb experiment only, so evidence from a subcutaneous plug or an in vitro construct did not raise the grade; independent donor-only vessels required a host-specific marker positively stained and negative in the donor-lined vessel; and a lipophilic membrane label alone was counted as donor-cell identification,^17^ with the stricter species- or genotype-specific rule reported as a sensitivity analysis.

Grading proceeded in three passes (extraction, adversarial verification instructed to refute over-claims, and a reverse-direction audit), and studies whose grade turned on figure detail were read directly by reviewers.

### Statistics

Effect sizes are Hedges’ g for the difference in limb perfusion between an intervention arm and the acellular control of its own experiment, computed with the bias correction for small samples.^18^ Where a study reported standard error, it was converted to standard deviation. Where the dispersion type was not stated, standard deviation was assumed and the assumption tested in sensitivity analysis. Where several intervention arms shared one control arm, the control sample size was used unsplit in the primary analysis and split across arms in sensitivity analysis.

The primary analysis pooled the comparison at each experiment’s primary timepoint (its latest, longest-follow-up perfusion measurement) in a three-level random-effects model (comparisons nested within studies)^19^ fitted by restricted maximum likelihood,^20^ with cluster-robust variance at the study level,^21^ using the bias-reduced CR2 estimator with Satterthwaite degrees of freedom^22, 23^ rather than the conventional CR1 estimator, which uses residual degrees of freedom and is known to understate uncertainty when clusters are few. Both estimators are reported in the Supplemental Data. A univariate random-effects model with the Hartung–Knapp adjustment is reported alongside as a confirmatory analysis;^24^ for that comparison the comparisons within each study were first aggregated to a single effect, accounting for the covariance induced by shared control arms, so that each study contributes as one independent unit^25^. Heterogeneity is summarized by I^2^, τ^2^, Cochran Q, and a 95% prediction interval. Uncertainty is reported throughout as 95% confidence intervals. Robustness of the pooled estimate to the choice of timepoint was assessed by repeating the primary analysis at fixed windows around day 14 and day 28, the latest-timepoint analysis remaining the primary one. Because a standardized mean difference can be inflated where the outcome is a bounded ratio, the perfusion effect was also expressed on the natural scale, as a log ratio of means across all comparisons and, within the subset reporting a 0–1 perfusion ratio, as a raw mean difference. Dependence among comparisons sharing a control arm was additionally modeled with a correlated-and-hierarchical-effects working model across a range of assumed within-study correlations. Small-study effects were assessed with a standard-error–based regression test and, because that test is confounded for standardized mean differences, with a sample-size–based regression test. The post hoc time-course meta-regression was repeated on the natural scale and under both cluster-robust estimators, and the control arm’s mean and coefficient of variation were regressed on time within studies to test whether spontaneous recovery of the control limb could inflate the standardized measure. Multilevel I^2^ components follow Cheung (2014)^26^; their formulas are given in the Supplement.

Eight moderators were prespecified: donor-cell fate grade, PSC source, delivery vehicle, timing of dosing, host immune status, host comorbidity, delivery route, and cell type. The delivery-route moderator has three levels: intramuscular injection, epimysial or subcutaneous implantation of a solid construct, and systemic administration (intra-arterial or intravenous). Route was assigned from the arms that contributed comparisons rather than from the study as a whole. Subgroup differences were tested by moderator within the model, and the estimate reported for each level is the corresponding cell mean of that same model, so that the variance components are shared across levels rather than re-estimated within each. P values were adjusted by the Holm–Bonferroni method across the prespecified set, which contributes seven tests because the cell-type moderator is not estimable.^27^ Three further moderators derived from the tracking domain are reported as post hoc, adjusted within their own family of three. The prespecified family is unaffected. For the prespecified secondary outcome of limb status, the event was defined as a preserved limb at the experiment’s final timepoint, using each study’s own definition (limb salvage, freedom from autoamputation, or freedom from limb loss), for the primary cell arm against the acellular control of the same experiment. Risk ratios were pooled in a DerSimonian–Laird random-effects model with a Hartung–Knapp interval as sensitivity, a 0.5 continuity correction for zero cells, and a risk-difference sensitivity analysis. Ordinal necrosis scores and behavioral tests, which are not commensurable across studies, are summarized without pooling.

Small-study effects were assessed by contour-enhanced funnel plot, Egger’s regression,^28^ Begg and Mazumdar’s rank correlation,^29^ Duval and Tweedie’s trim-and-fill using both L0 and R0 estimators,^30^ and the Ioannidis–Trikalinos excess-significance test.^31^ Analyses used R 4.5.2 with the metafor package (version 5.0.1)^20^ and the clubSandwich package (version 0.7.0).^23^ All analysis code is available in the repository, and all reported statistical results, tables, and figures were generated directly from the extracted dataset using this code, rather than manually transcribed.

## Results

### Most Included Studies Contributed Primary-Timepoint Comparisons to the Pooled Estimate

The searches returned 710 records: 105 from PubMed under the field-unrestricted strategy, 9 from the registered field-restricted supplement, 98 from Web of Science, 200 from Europe PMC, and 298 from Embase. We removed 272 duplicates and screened 438 records, of which 310 were excluded. Full texts were sought for 128 records, and 55 were excluded at full text.

Seventy-three studies met the criteria at full-text assessment. One study was excluded during data extraction because it reports no limb perfusion outcome: laser Doppler was used only to confirm that ischemia had been induced, the quantity called “blood flow” is FITC-dextran within a subcutaneous Matrigel plug, and the primary comparison is cell versus cell with no acellular arm.^32^ One pair of reports describes a single cohort, with the same ligation “performed as described previously”, the same dose of 6×10^5^ blast cells, the same cell-free medium control, and the same 4-week endpoint. That pair is counted as one study, with the effect estimate attributed to the earlier report and the later report contributing only to the donor-cell fate record.^13, 14^ Four further studies, together with the separately allocated mesenchymal stromal arms of two others, were excluded as mesenchymal stromal (non-vascular lineage) products (Methods).

The review includes 68 studies described in 69 reports (**Figure 1**). Fifty contribute at least one comparison. Eighteen contribute none, for the following reasons: no perfusion outcome (4); no acellular control (7); a figure that is unobtainable or not quantitative (4); a control arm reported as exactly zero with no dispersion, so that no standardized mean difference exists (1); perfusion shown only as a dot plot, interquartile range, or violin without a usable mean and dispersion (1); a perfusion experiment performed with primary rather than pluripotent stem cell-derived endothelial cells (1). These 18 remain in the review and in the narrative synthesis (**Supplemental Table S1**).

**Figure 1.**
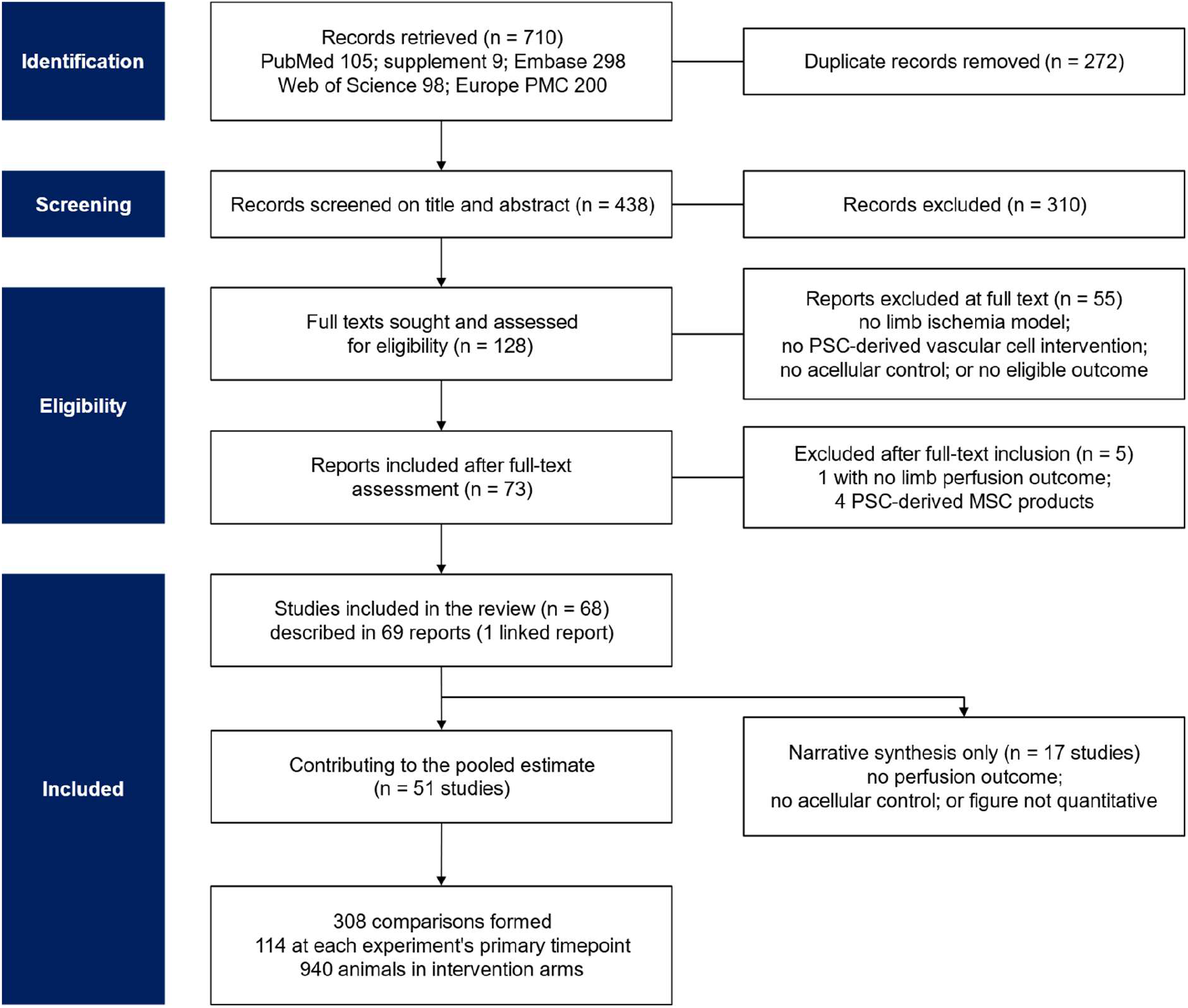
Study selection (PRISMA 2020). Records were retrieved from PubMed with a field-unrestricted strategy and with the registered field-restricted supplement, from Web of Science Core Collection, from Europe PMC, and from Embase. One study was excluded after full-text inclusion, at the data-extraction stage, because it reports no limb perfusion outcome and has no acellular control. A further 4 studies were excluded as mesenchymal stromal (non-vascular lineage) products (Methods). One further report describes the same animals as an included study and is counted as a linked report rather than as a separate study. Every count in the diagram is computed from the screening files rather than transcribed. The 68 studies that met inclusion comprise 50 that contribute at least one quantitative perfusion comparison to the pooled analysis and 18 carried in narrative synthesis only. PRISMA indicates Preferred Reporting Items for Systematic Reviews and Meta-Analyses.

### Included Studies Were Small and Heterogeneous and Dominated by Healthy Young Mice

To describe the evidence base, we summarized the design of each included study. The 68 studies span 2007 to 2026, with 44 published in 2015 or later (**Table 1**; **Supplemental Table S1**).^13, 14, 17, 33–98^ All 68 used mice; no study in a large animal met the eligibility criteria. Fifty-five used immunodeficient hosts, 9 immunocompetent or humanized hosts, and 4 more than one host within the same paper. Sixty used healthy young animals. Only 8 used a comorbid or aged model, although the target indication is CLTI in older patients with diabetes and atherosclerosis. Delivery was by intramuscular injection in 58 studies. These data show that the literature is dominated by healthy young mice rather than by comorbid or aged models.

**Table 1.**
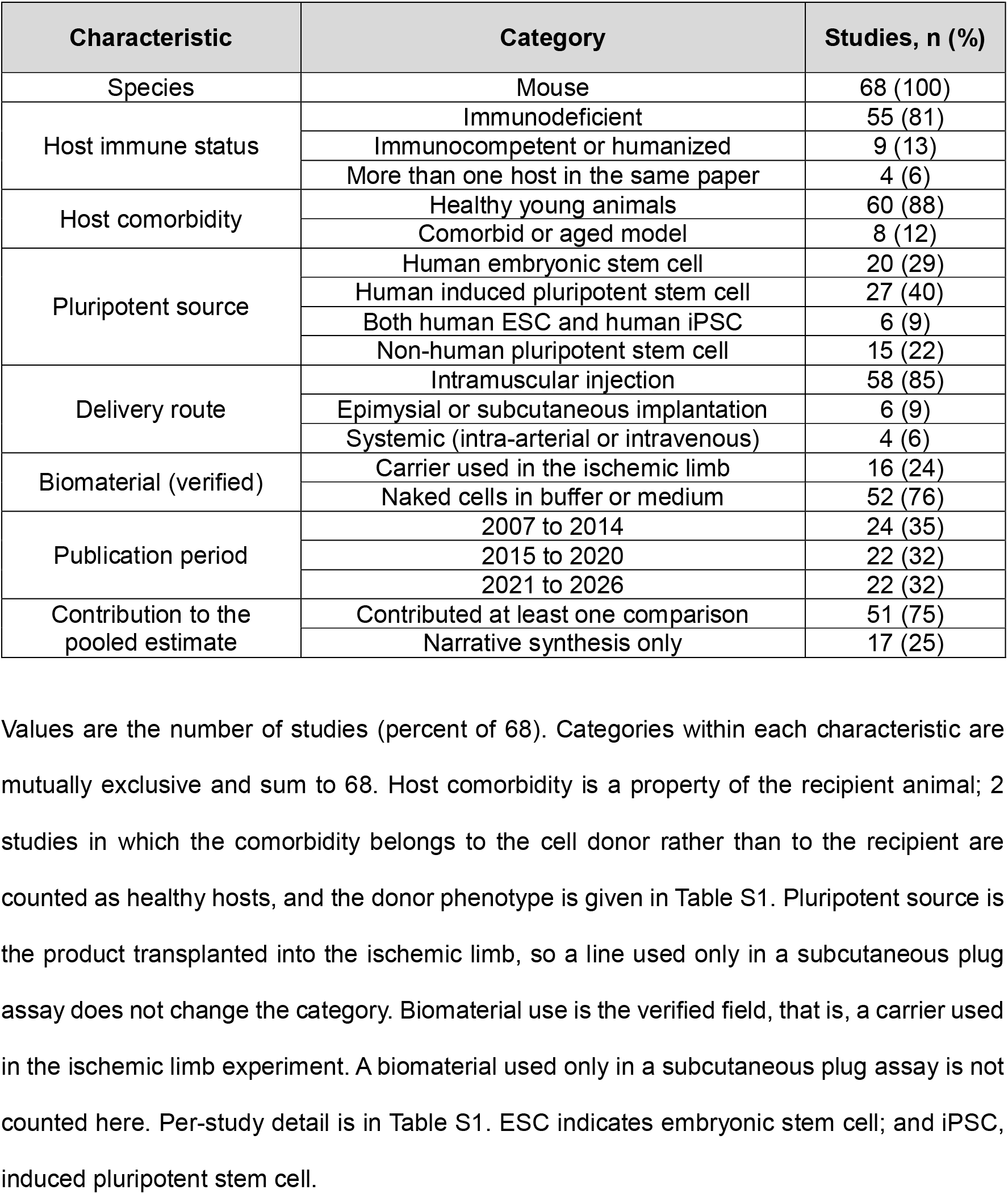
Characteristics of the 68 included studies.

| Characteristic | Category | Studies, n (%) |
| --- | --- | --- |
| Species | Mouse | 68 (100) |
| Host immune status | Immunodeficient | 55 (81) |
|  | Immunocompetent or humanized | 9 (13) |
|  | More than one host in the same paper | 4 (6) |
| Host comorbidity | Healthy young animals | 60 (88) |
|  | Comorbid or aged model | 8 (12) |
| Pluripotent source | Human embryonic stem cell | 20 (29) |
|  | Human induced pluripotent stem cell | 27 (40) |
|  | Both human ESC and human iPSC | 6 (9) |
|  | Non-human pluripotent stem cell | 15 (22) |
| Delivery route | Intramuscular injection | 58 (85) |
|  | Epimysial or subcutaneous implantation | 6 (9) |
|  | Systemic (intra-arterial or intravenous) | 4 (6) |
| Biomaterial (verified) | Carrier used in the ischemic limb | 16 (24) |
|  | Naked cells in buffer or medium | 52 (76) |
| Publication period | 2007 to 2014 | 24 (35) |
|  | 2015 to 2020 | 22 (32) |
|  | 2021 to 2026 | 22 (32) |
| Contribution to the pooled estimate | Contributed at least one comparison | 51 (75) |
|  | Narrative synthesis only | 17 (25) |

The cell products are heterogeneous even within nominally identical categories: endothelial and endothelial-like cells predominate, alongside vascular progenitors, hemangioblasts, pericytes, smooth muscle cells, vascular organoids, and co-delivered endothelial-plus-mural combinations. Follow-up was 14 days in the largest group and 28 days in the next. Four studies followed animals for 64 days or more. These data indicate that neither the cell product nor the duration of follow-up is standardized across this literature.

### Risk of Bias Was Unclear Across Most SYRCLE Domains Because of Non-Reporting

To assess internal validity, we applied the SYRCLE risk-of-bias tool to each included study. Across the ten SYRCLE domains (**Figure 2A**; **Supplemental Table S2**), no study described allocation concealment, none allowed selective outcome reporting to be judged against a protocol, and none described blinding of caregivers. Sequence generation was the best-reported domain, and even there only 28 of 68 studies were low risk.

**Figure 2.**
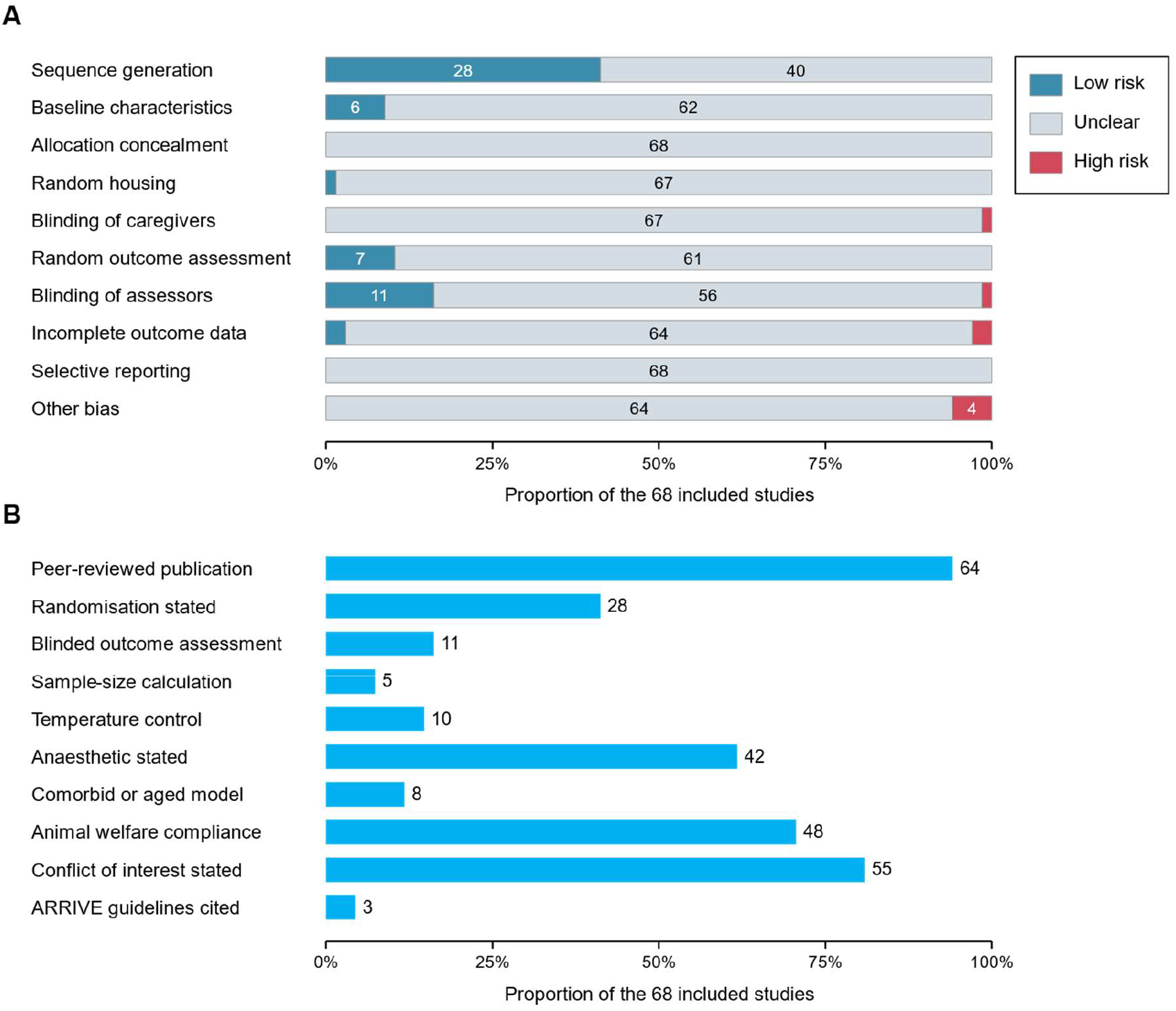
Risk of bias and reporting quality. (A) SYRCLE judgments across the 10 domains, shown as the proportion of the 68 included studies at each judgment, with study counts printed in the bars. (B) The proportion of studies meeting each item of the adapted CAMARADES checklist. Unclear judgments dominate because the information was not reported, not because a flaw was demonstrated. All 68 included studies are scored here, including the 17 carried in narrative synthesis only. CAMARADES indicates Collaborative Approach to Meta-Analysis and Review of Animal Data in Experimental Studies; and SYRCLE, Systematic Review Centre for Laboratory Animal Experimentation.

The CAMARADES median score was 4 of 10 (range 0–9; **Figure 2B**). Randomization was described in 28 studies, blinded outcome assessment in 11, a sample-size calculation in 5, and the ARRIVE guidelines^99^ were cited by 3. These gaps bear on interpretation and not only on completeness of reporting. Laser Doppler perfusion imaging is an operator-dependent measurement of a ratio, and fewer than one study in six describes blinded assessment. These results indicate that a literature that does not blind the measurement of its own primary outcome cannot exclude expectation effects.

### Cell Therapy Increased Limb Perfusion by a Large Amount

To estimate the effect of cell therapy on limb perfusion, we pooled the primary-timepoint comparisons from the contributing studies. Fifty-one studies contributed 308 comparisons, of which 114 at each experiment’s primary timepoint entered the primary analysis, representing 940 animals in intervention arms and 573 in acellular control arms (1,513 in total). Cell therapy increased limb perfusion relative to acellular control: Hedges’ g 2.26 (95% CI 1.72–2.79; P=4.1×10-11) in the three-level model with cluster-robust variance (**Figure 3; Supplemental Figure S1**). The univariate model with the Hartung–Knapp adjustment gave g 2.14 (95% CI 1.72–2.57). Aggregating each study to a single effect and applying the same adjustment, the confirmatory estimate was g 1.71 (95% CI 1.10–2.32). On the natural scale, treated limbs recovered about twice the perfusion of their own controls (ratio of means 1.96, 95% CI 1.66– 2.33); within the studies reporting a bounded 0–1 perfusion ratio the raw mean difference was 0.26 (95% CI 0.21–0.31). The pooled estimate was similar at fixed follow-up windows (day 14, g 1.85 [95% CI 1.45–2.25]; day 28, g 2.19 [1.66–2.73]), and a correlated-and-hierarchical-effects model gave g 1.99 to 2.12 across assumed within-study correlations, so the direction of benefit did not depend on the timepoint chosen or on the handling of shared controls, even though the standardized estimate was larger at the later window (**Supplemental Table S3**).

**Figure 3.**
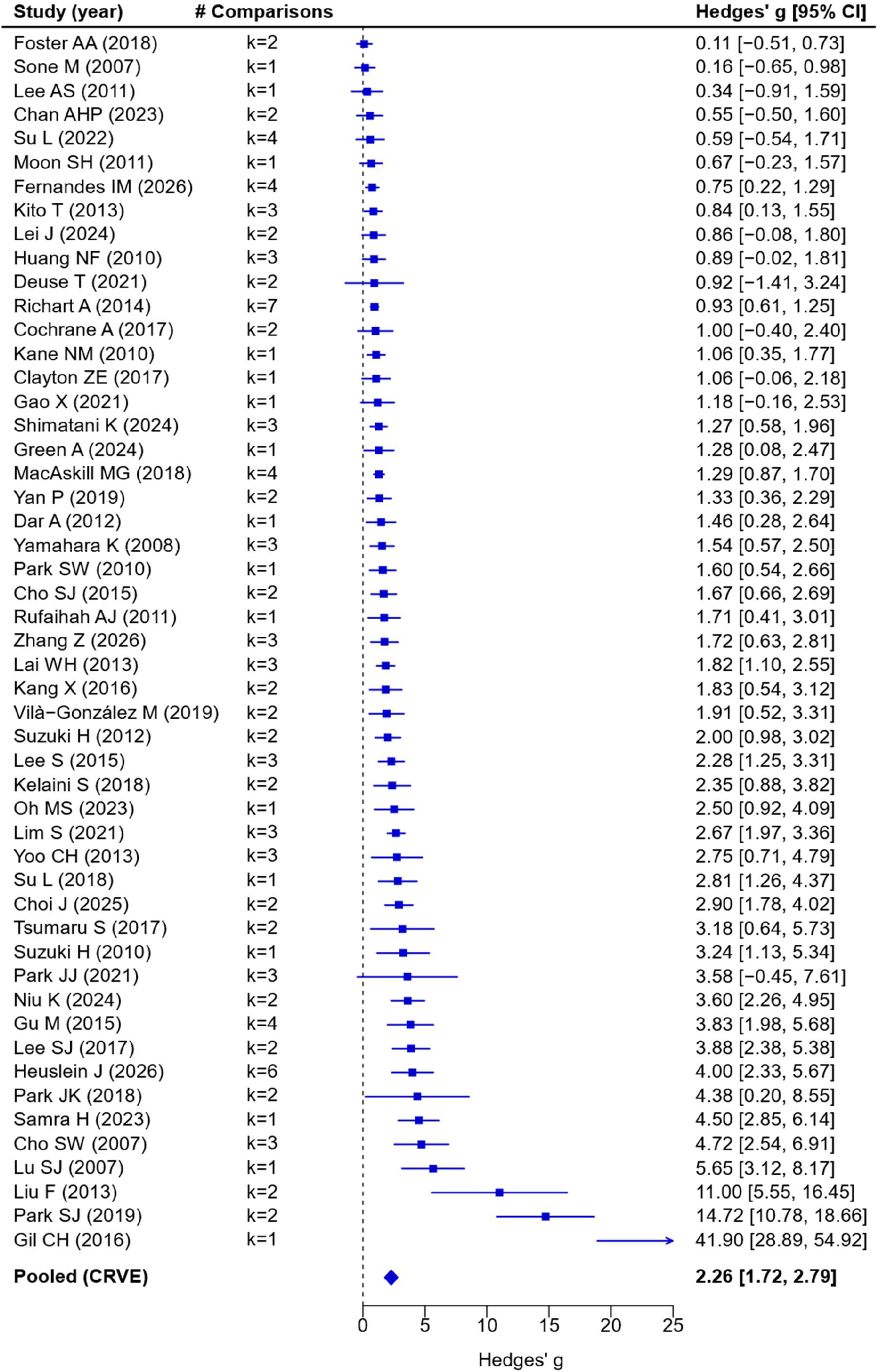
Perfusion recovery after PSC-derived vascular cell therapy. The plot is built from the 50 of 68 included studies that provide at least one quantitative perfusion comparison against an acellular control (114 comparisons in total). The other 18 studies report no poolable perfusion outcome and are carried in narrative synthesis only. There is one row per contributing study, ordered by effect size and labeled by first author and year. Where a study contributes several intervention arms, they are pooled first in a three-level model, so no study is counted more than once. k is the number of comparisons behind that row. The diamond is the prespecified three-level model with cluster-robust variance (CRVE). Positive values favor cell therapy. The horizontal axis is clipped at 20; one study whose interval runs beyond it is drawn as an arrow and its value is given in full in the estimate column. Figure S1 shows the same evidence as all 99 individual comparisons. CI indicates confidence interval; CRVE, cluster-robust variance estimation; and PSC, pluripotent stem cell.

The magnitude of this estimate should be interpreted with care. A standardized mean difference of 2.5 corresponds to the average treated animal exceeding roughly 99% of control animals. Effects of that size are not seen in the clinical literature on cell therapy for limb ischemia,^3, 4^ and in preclinical work an estimate of this size usually reflects unresolved variance and selective reporting in addition to a true treatment effect.

One contributor is specific to the outcome measure: most studies report a bounded ratio of ischemic to contralateral perfusion, whose compressed control-arm variance inflates a standardized mean difference independently of any biology and also distorts funnel-based tests of small-study effects. The most extreme instance is one comparison whose reported standard deviations at 4 weeks are about 2% of the arm means, giving a standardized mean difference of 41.9; it holds 0.035% of the weight in the pooled model, and removing the study moves the estimate only from 2.26 to 2.19, but it shows plainly how a bounded ratio with a near-constant control arm behaves on the standardized scale,100 so a modality-restricted sensitivity analysis is reported and the bias analyses are interpreted cautiously.

### Heterogeneity Was Very High and the Prediction Interval Spanned Harm to Benefit

Heterogeneity in the primary analysis was high, in the considerable range on the Higgins scale101: I2=87.5%, τ2=2.59, and Cochran Q=591.0 on 113 df (P<2×10-16) in the univariate model. In the three-level model the variance partitioned as 71.9% between studies and 18.6% within studies, 90.5% in total, so most inconsistency lies between laboratories rather than between arms of one experiment. The 95% prediction interval runs from −1.47 to 5.98; the corresponding interval from the univariate model is −1.07 to 5.36. A new study in this literature could plausibly find substantial harm, no effect, or an enormous benefit. These results indicate that the direction of benefit is consistent across this literature, but that a prediction interval of this width does not allow the effect size of a future study to be anticipated.

### Limb Preservation Favored Cell Therapy in the Minority of Studies Reporting It

Twenty-five of 68 studies (37%) report any functional or limb-status outcome and 43 report none (**Figure 4**). Eleven studies provide countable limb-status events for the prespecified secondary outcome, two of them with control counts read from published stacked-bar figures. A twelfth reports only a group-level statement and enters sensitivity analysis only. Seven of the eleven control arms recorded no events, so the continuity correction applied to zero cells pulls the pooled ratio below the ratio of the crude totals. Across the eleven, 65 of 109 cell-treated animals versus 7 of 108 acellular-control animals kept a preserved limb at the final timepoint, a pooled risk ratio of 4.88 (95% CI 2.28–10.45; I^2^=28%; risk difference 0.50, 95% CI 0.34–0.67) (**Figure 4A**). The estimate warrants the same caution as the perfusion estimate. Seven of the eleven control arms had zero preserved limbs, so the ratio is driven by extreme event rates in autoamputation-prone acute models, and leave-one-out estimates range from 3.75 to 7.70. Eight further studies report only an ordinal necrosis or tissue-damage score, each favoring the cell arm, and two only a narrative claim. Behavioral or functional testing, the outcome closest to what a patient would notice, appears in four studies (6%): Tarlov and Faber scores improved significantly in one, quantified gait improved in one, a limb-function score improved in one, and treadmill distance showed a non-significant trend in one. **Figure 4B** shows three of them because the fourth also provides countable limb-status events, and each study is displayed once at its most informative level. These results indicate that the direction matches the perfusion analysis. Limb preservation, the outcome closest to the clinical endpoint, is counted in about one study in six, so the evidence for it is smaller in volume than the evidence for perfusion and consistent with it.

**Figure 4.**
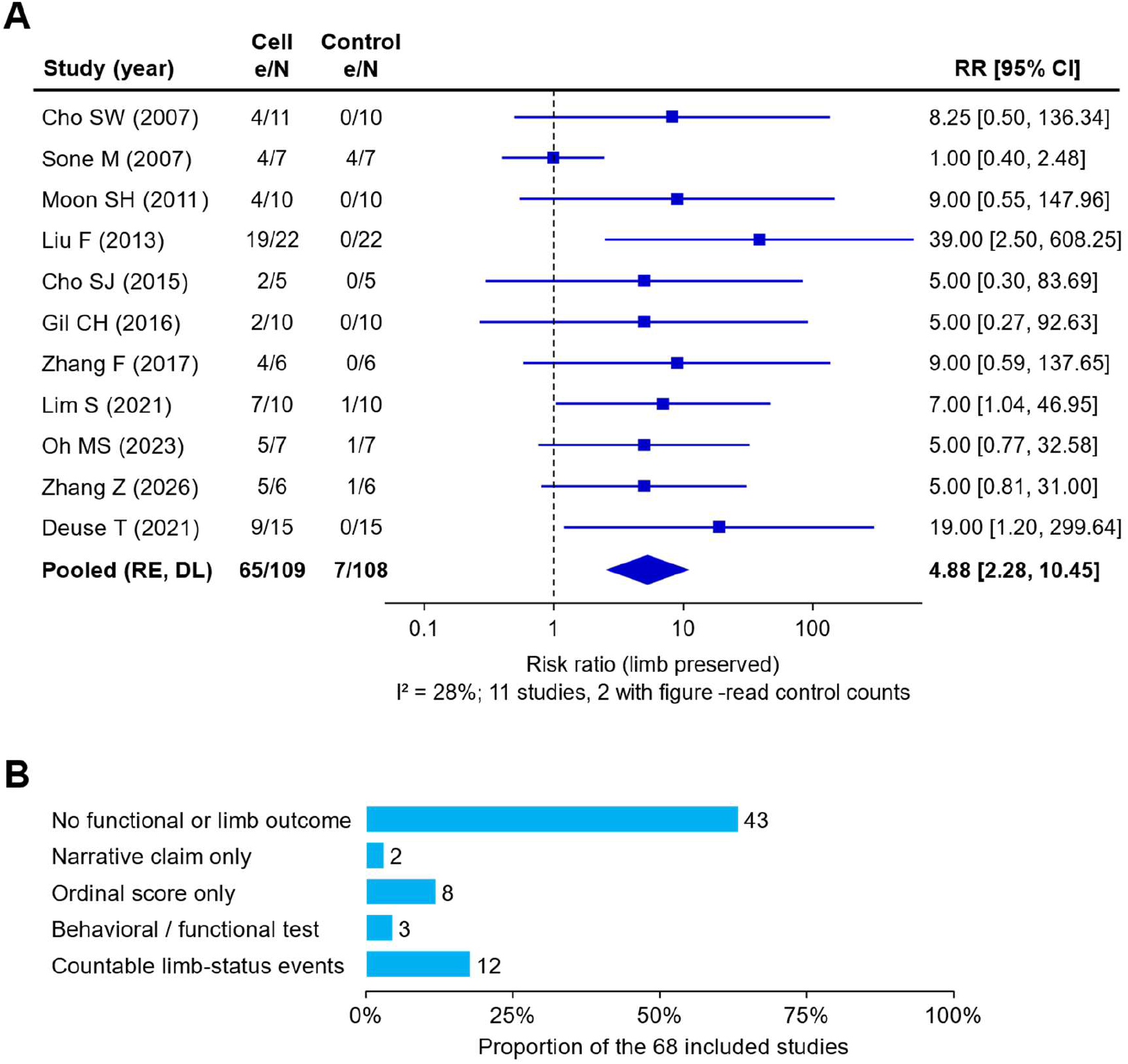
Functional and limb-status outcomes across the 68 included studies. (**A**) Risk of a preserved limb at each experiment’s final timepoint in the 11 studies contributing countable limb-status events to the pooled estimate. The event is defined per study as limb salvage, freedom from autoamputation, or freedom from limb loss, for the primary cell arm against the acellular control of its own experiment. The model is a random-effects model with a 0.5 continuity correction for zero cells, and the totals are 65/109 versus 7/108. For 2 studies the control counts were read from published stacked-bar figures. (**B**) Reporting of functional outcomes across all 68 studies, each study counted once at its most informative level. Per-study event definitions, counts, and source quotations are deposited with the analysis code.

### No Prespecified Moderator Including Donor-Cell Fate Grade Explained the Heterogeneity

To determine whether the heterogeneity was explained by study characteristics, we tested the prespecified moderators. No prespecified subgroup difference survived Holm–Bonferroni adjustment (**Figure 5**). The pooled estimate and comparison count (k) within each stratum are shown there. No prespecified moderator reached a raw subgroup P value below 0.05 (raw P 0.23 to 0.92), and every one had an adjusted P of 1.00. The widest spread was for delivery route (intramuscular injection g 2.51, epimysial or subcutaneous implantation 1.49, systemic 1.09; raw P=0.23), but each non-intramuscular stratum draws on only 3 studies, and the cluster-robust denominator degrees of freedom (2.4) fall below the value at which this test is dependable. Moderator strata are counted in comparisons (k), whereas **Table 1** counts studies (n), because a study may contribute several comparisons.

**Figure 5.**
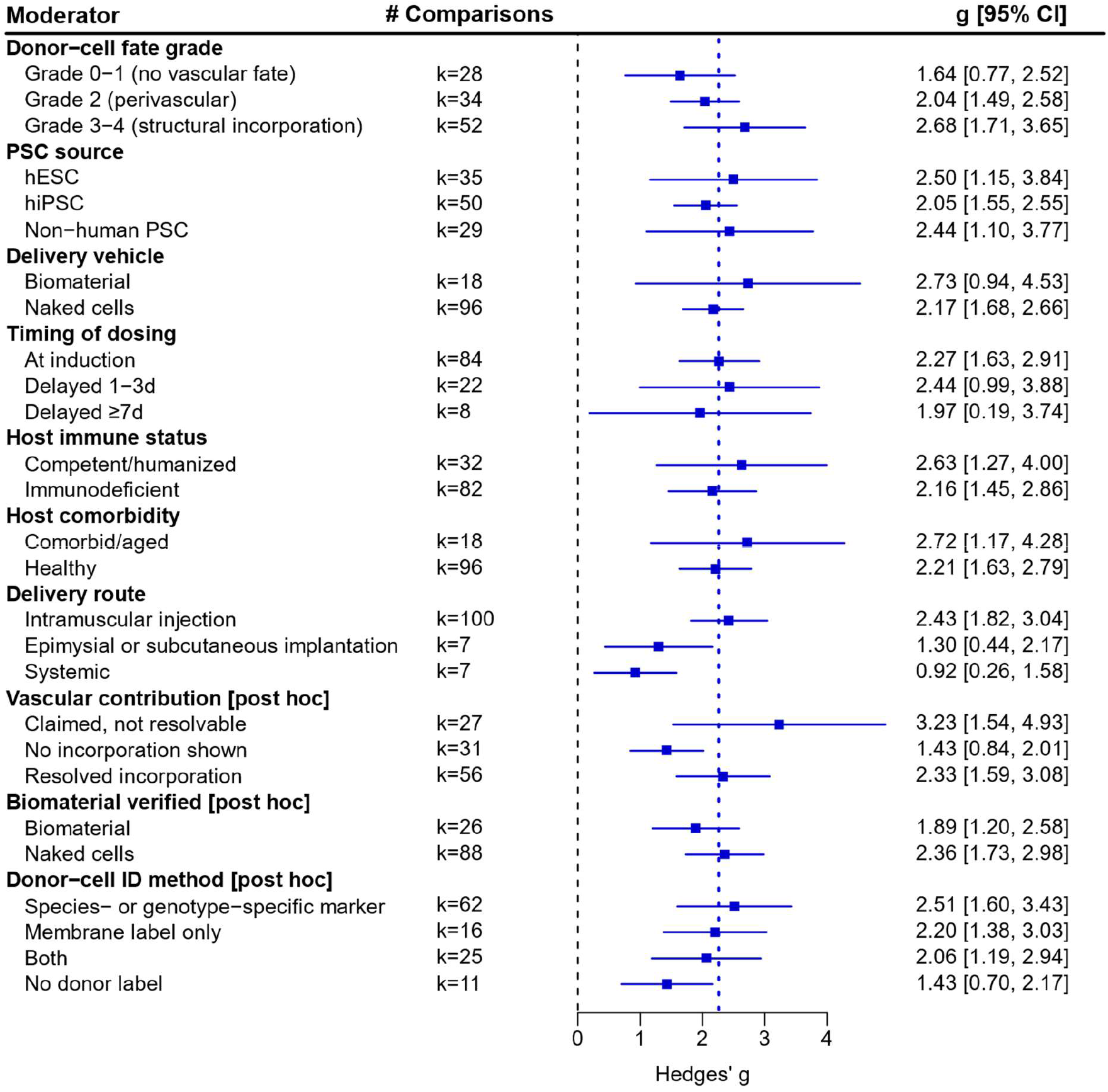
Effect by every moderator examined. Each row is the pooled estimate within one stratum from a three-level model with cluster-robust variance. k is the number of comparisons. The strata partition the same 114 comparisons from 51 studies shown in Figure 3. The dotted vertical line is the overall pooled estimate and the dashed line is no effect. The 3 post hoc moderators are labeled as such; they stratify the same 114 comparisons. Each level is the cell mean of a single three-level model for that moderator, so the variance components are shared across its levels. The cell-type moderator as recorded is not shown because it is not estimable; a post hoc collapsed version is reported in the text. The epimysial or subcutaneous implantation stratum holds 7 comparisons from 3 studies: a hydrogel patch and a cell sheet placed on the ischemic muscle, and one capsule implanted at an ectopic subcutaneous site away from the limb. No moderator difference survived Holm–Bonferroni adjustment within its family.

The fate-grade moderator was the review’s primary mechanistic hypothesis, and it is null (P=0.30). The five grades were collapsed to three strata for estimation (grades 0–1, no vascular fate; grade 2, perivascular; grades 3–4, structural incorporation), and the point estimates are 1.78, 2.15 and 2.72, so they do not order with the grade, and the intervals overlap almost completely. This is the moderator with the most adjudication behind it: every study was graded, re-graded in the opposite direction, and, where figures were decisive, read directly by reviewers. The null result is therefore not an artifact of loose grading. The cell-type moderator is not estimable, because 42 distinct product descriptions were recorded across the 51 contributing studies and 15 appear once. Collapsing these descriptions post hoc into 5 lineage groups leaves the moderator null (endothelial or endothelial-like 2.17, 57 comparisons; vascular progenitor 2.10, 19; endothelial plus mural co-delivered 2.08, 16; mural cell 1.27, 5; hemangioblast 9.42, 2; QM P=0.20), although the two smallest groups carry intervals too wide to interpret. These data indicate that the field has not converged on a comparable product.

Two moderators were assigned per experiment rather than per study, because six papers run the same cells in more than one host and two run diabetic and non-diabetic experiments side by side. Host immune status was read from the strain statement of each experiment, which brought previously unassigned comparisons into the analysis. Host comorbidity was assigned the same way, and two studies whose recorded comorbidity belongs to the cell donor rather than the recipient were reclassified as healthy hosts. Neither correction rescued either moderator. Both became more clearly null.

We then performed continuous meta-regression, which showed no association with publication year (slope −0.011, P=0.82; the same trend is shown cumulatively in **Supplemental Figure S2**), CAMARADES score (slope 0.002, P=0.99), or timepoint (slope 0.026, P=0.44).

### Donor Cells in Vessel Walls Were Predominantly Mosaic With Host Cells

To determine the fate of donor cells, we recorded the vascular contribution reported by each study. Fifty-seven of 68 studies (84%) attempted to detect donor cells in host tissue, and 34 (50%) counted them. The seven-level vascular-contribution breakdown across the 68 studies is shown in **Figure 6A**, with per-study records in **Supplemental Table S4**. Thirty-five studies of 68 resolved a donor cell to a vessel wall, and 32 of those demonstrated a wall that also contained host cells. On the prespecified five-level fate grade the 68 studies distribute as 9 at grade 0, 12 at grade 1, 14 at grade 2, 30 at grade 3, and 3 at grade 4. The dominant biology in this literature is therefore not the construction of new donor vessels but the insertion of donor cells into vessels that host cells are also building. Five studies demonstrated a vessel composed of donor cells alone, three exclusively^35, 38, 62^ and two alongside mosaic vessels in the same paper.^91, 94^ Counts are of studies, and the two studies showing both patterns are counted in both the mosaic and the donor-only totals, which is why those two totals sum to more than the 35 studies that resolved a donor cell to a vessel wall. In one, the donor-only vessels were nonetheless shown to anastomose with the host circulation by perfusion, intraluminal erythrocytes, and lectin.^38^ Anastomosis and wall chimerism are separate axes.

**Figure 6.**
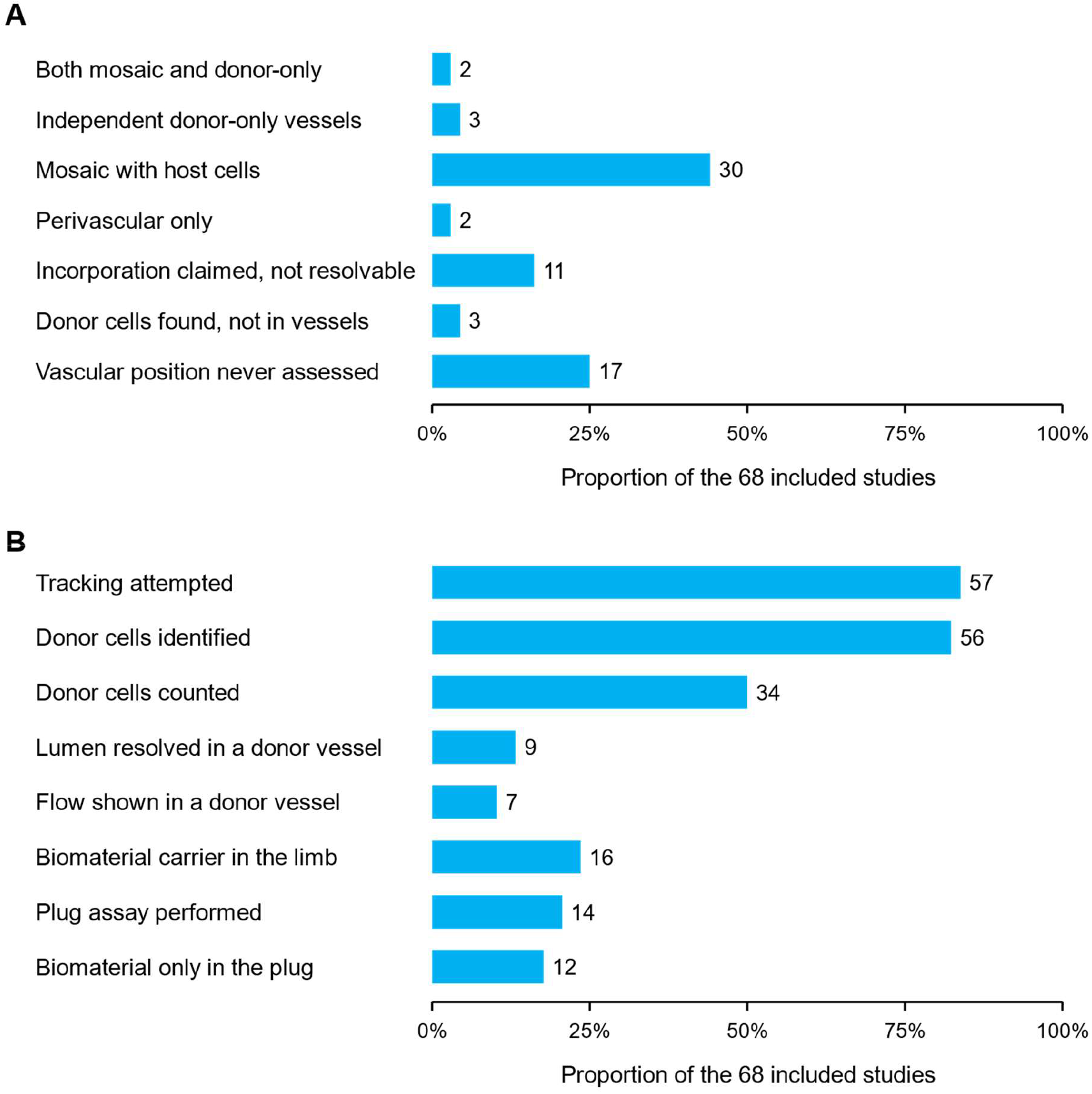
Donor-cell tracking across all 68 included studies, assessed independently of whether a study contributes a perfusion comparison. (**A**) The seven-level vascular-contribution ordinal. (**B**) The remaining extraction criteria, as the proportion of the 68 included studies meeting each, with the count printed at the end of each bar. Lumen resolved in a donor vessel and flow shown in a donor vessel fall far below tracking attempted: membership of a vessel wall is not the same as participation in a perfused conduit.

The criterion met by these 32 studies is less stringent than demonstration of a perfused donor-containing vessel. A patent lumen in a donor-containing vessel was resolved in 9 studies, and flow through such a vessel, by systemically administered lectin, dextran, or intraluminal erythrocytes, was resolved in 7 (**Figure 6B**). These results indicate that membership of a vessel wall is not the same as participation in a perfused conduit.

Two of the eleven studies in this stratum sit at *incorporation claimed, not resolvable* because of the review’s own rule rather than any weakness of the papers. In both, a donor cell demonstrably occupies the wall of a vessel with a patent lumen: a Y-chromosome-positive nucleus within an anti-human-CD31-positive wall in one,^55^ and human PECAM-positive microvessels with resolved lumens in the other.^53^ Neither stains a host-specific marker anywhere, so *independent* cannot be awarded, and without a host co-stain neither can *mosaic*.

Seventeen studies never asked where donor cells sat relative to vessels. Every one of them was examined twice in opposite directions, so this is a finding about the field rather than a limitation of extraction. All seventeen fall into three designs. First, donor cells were followed by whole-animal bioluminescence for survival only. Second, limb sections were stained with host-specific or pan-species markers alone. Third, donor-cell evidence was generated in a plug assay or in vitro and never applied to the limb.

The scope rule mattered: a subcutaneous plug assay in a healthy animal shows that the cells can form vessels, not that they did so in the ischemic limb, and grading only the therapeutic experiment moved several studies down.^74, 79, 89^

The same distinction corrects the biomaterial count: 16 studies used a biomaterial as a therapeutic carrier in the limb, whereas a rule based on the reported delivery vehicle would have classified 19, so the classification was taken from the therapeutic experiment.

### Most Studies Labeled Donor Cells but Some Lacked Transfer Controls

Fifty-six studies applied at least one donor-cell label, and 47 of those used a species- or genotype-specific marker. Nineteen studies used an exogenous label. Ten paired it with an orthogonal species-specific or genetic marker, which disposes of the transfer concern directly: a transferred membrane label cannot confer a human nucleus, human CD31, or reporter expression. One study went further and quantified ^14^C by accelerator mass spectrometry in ischemic muscle and five distant organs at eight timepoints to day 90 and measured ^14^C excretion, so that the fate of the label was measured rather than assumed.^45^

Nine studies are based on an exogenous label alone, and not one performed a dedicated transfer control: no macrophage co-stain, no dead-cell arm, and no dye-only arm. For five of the nine a species-specific marker was impossible by design, because the graft was syngeneic mouse-into-mouse. For the remaining four a species-specific reagent was available and was not applied to the limb. The limits of in vivo cell tracking are not specific to this literature: no single label distinguishes a surviving donor cell from label transferred to a host phagocyte, which is why an orthogonal, species- or genotype-specific marker is the safeguard rather than the dye itself.^100^

### Reverse-Direction Auditing Raised Resolved Vascular-Contribution Calls and Changed Many Gradings

Grading was performed by an extractor and verified adversarially by an independent reviewer instructed to refute over-claims; because that design can only detect false positives, every conservative call was then re-audited in the opposite direction, and studies whose grade turned on figure detail were read directly by reviewers. Studies whose vascular contribution was resolved to a vessel wall rose from 6 at first grading to 32 of the 63 audited studies, and 30 of the 63 final calls differ from the call made at first grading; one conference abstract remains permanently unresolvable because it does not contain the confocal images a decision would need.^98^ The five studies added by the Embase search were graded once against the same rubric and three resolved a donor cell to a vessel wall, so 35 of the 68 studies reached a resolved call; every call is retained in the deposited dataset with its confidence rating, reasoning, decisive quotation, and source location.

### The Largest Effect Came From Studies With Unresolvable Incorporation Claims

The three moderators derived from the tracking domain (the method each study used to identify donor cells, the collapsed vascular-contribution stratum, and the verified biomaterial field) are post hoc and were adjusted within their own family. None survives adjustment, and all are hypothesis-generating. The pooled estimate within each post hoc stratum is shown in **Figure 5**. Effects were ordered by the specificity of the donor-cell identification method, from a species- or genotype-specific marker (g 2.67) through a membrane label alone or both marker types (2.16 and 2.11) to no donor label (1.53), but the gradient was not significant (raw P=0.43, adjusted P=0.43).

The vascular-contribution result runs opposite to the prevailing mechanistic account. The pooled effect is largest in studies whose incorporation claims their images cannot resolve (g 3.23), lower in studies that resolved incorporation (g 2.33), and lower again in studies showing no incorporation (g 1.43). Isolating this claimed-but-not-resolvable stratum reverses the direction of the prespecified fate grade, in which the structural-incorporation stratum carried the largest estimate. If donor-cell incorporation drove functional benefit the ordering would be reversed. The contrast became larger when reviewers read the disputed figures. Re-reading moved comparisons out of the unresolvable stratum into the other two, leaving 27 comparisons from 11 studies whose incorporation claims their images still cannot resolve, and those remaining have the largest effects.

The registered vehicle rule gave biomaterial g 2.73 versus 2.17 for naked cells (P=0.56), whereas the verified field reversed the direction (1.89 versus 2.36, P=0.30); neither is causal, but the contrast shows that a moderator built from the reported delivery vehicle can be directionally wrong.

### The Apparent Widening With Longer Follow-Up Was Not Robust to Scale or Variance Estimator

The primary analysis uses one timepoint per experiment and sets 194 of 308 comparisons aside. Using all 308 in a three-level meta-regression, g rises by 0.039 per day, but the slope is significant only under the CR1 estimator (95% CI 0.008–0.070, P=0.016) and not under the bias-reduced CR2 estimator (95% CI −0.025 to 0.102, P=0.14). The two estimators are also tested differently, CR1 with residual and CR2 with Satterthwaite degrees of freedom; applying Satterthwaite degrees of freedom to CR1 as well leaves the slope non-significant (P=0.07), so this contrast reflects the treatment of degrees of freedom as much as the choice of estimator (**Supplemental Figure S3**). Restricted to the 99 primary-timepoint comparisons the slope falls to 0.026 and is not significant. The widening is also scale-dependent: on the natural scale it is attenuated and no longer significant (log ratio of means 0.0059 per day, 95% CI −0.0015 to 0.0134, P=0.09), and the fixed windows diverge in the same direction, the standardized and natural-scale estimates agreeing at day 14 (g 1.89; ratio of means 1.84) but not at day 28 (g 2.28; ratio of means 1.98). A measurement explanation fits this pattern. Control animals recover spontaneously in this model, and within studies the control mean rises with time (P=7×10⁻⁵) while the control coefficient of variation falls (P=5×10-8), which inflates a standardized mean difference at later timepoints independently of any biology. A widening effect is also not the shape simple acceleration of healing would produce, and laboratories may have continued measuring precisely where an effect was still visible. The apparent time course is therefore hypothesis-generating rather than established.

### No Dose–Response Was Detectable Across the Range of Cell Doses

Cell dose was extracted but not prespecified as a moderator. It is analyzed here because a product said to work by forming vessels might be expected to show dose dependence. Dose is free text in the source papers and was parsed once into a deposited file, with the rule that produced each value recorded. Of the 68 studies, 60 report a dose expressible in cells, spanning 4×103 to 1×107 cells and covering 105 of the 114 comparisons. Hedges’ g changed by +0.12 per ten-fold increase in dose (95% CI −1.28 to 1.02, P=0.77). No dose–response is detectable across nearly four orders of magnitude. This is a further prediction of the mechanistic account that the data do not show, although dose is compared across studies rather than within them, so it is confounded with cell product, model, and host, and the absence of a gradient is weaker evidence than a within-study dose escalation would provide.

### Small-Study Effects Persisted After Accounting for Clustering Within Studies

Because the 114 comparisons are nested within 51 studies (**Supplemental Figure S4**), and both Egger’s and Begg’s tests assume independent units, the tests are reported here on one effect per study, each study’s comparisons having been pooled first. The all-comparison versions are retained as a sensitivity check.

On the 50 study-level effects, Egger’s regression gave t=6.25 on 49 df (P=9×10-8) and Begg and Mazumdar’s rank correlation Kendall’s τ=0.48 (P=3.0×10-7). Using all 114 comparisons the same tests give t=11.16 on 97 df and τ=0.57, so clustering inflates the test statistics without changing the conclusion. A regression test that uses each study’s sample size in place of its standard error, avoiding the mechanical correlation between a standardized mean difference and its standard error, found no significant asymmetry (P=0.80) and a limit estimate of 2.61 (95% CI −1.07 to 6.28); the standard-error–based asymmetry is therefore at least partly an artifact of the effect measure rather than evidence of suppression (**Supplemental Table S5**).

The limit estimate does change, and the difference is material. Extrapolating the regression to zero standard error gives b=−0.22 (95% CI −0.87 to 0.43) on study-level effects, against −1.22 (−1.76 to −0.69) on the clustered comparisons. The clustered value would appear to imply that an infinitely precise study reports harm. That implication does not hold: once the units are independent, the extrapolated intercept is indistinguishable from zero. The defensible statement is that the apparent benefit shrinks toward nothing as precision increases, not that it reverses. Limit estimates of this kind are themselves biased under heterogeneity and are not offered as bias-corrected effects.

Trim-and-fill imputed 13 studies under L0, moving the estimate to 1.42. Under R0 it imputed 20 studies at the study level, moving the estimate from 2.00 to 1.15 (95% CI 0.48– 1.81). On all comparisons it imputed 22 and moved 2.14 to 1.47 (0.97–1.97). Both estimators impute, which is a stronger indication of funnel asymmetry than the L0 estimator alone would give, and both move the estimate in the same direction. Two caveats belong with these numbers. First, the method assumes funnel asymmetry arises from suppression rather than from genuine between-study variance and performs poorly at this level of heterogeneity, so it is illustrative rather than corrective. Second, because it cannot be applied to a multilevel model, its baseline is the univariate estimate and not the headline three-level estimate of 2.26; that univariate baseline of 2.00 aggregates each study without modeling the covariance induced by shared control arms, which is why it sits above the covariance-aware confirmatory estimate of 1.71.

The excess-significance test found 82 individually significant comparisons against 93.7 expected (χ2=8.3, P=0.004). The deficit is statistically significant, but it runs opposite to the pattern selective reporting produces, which is an excess of significant results rather than a shortage. Two features limit what the test can say here. Its expected count is computed at the pooled estimate, so if that estimate is inflated by the small-study effects the other analyses detect, the expected count is inflated with it and a deficit follows arithmetically; and with a pooled g of this size the power calculation saturates near 1, leaving little room for the test to discriminate. The result is therefore consistent with an overstated pooled magnitude and uninformative about suppression, and we do not read it as evidence either for or against selective reporting.

### Formal GRADE Was Not Applied and Certainty Is Low

GRADE is built for human clinical evidence, and its domains do not transfer cleanly to animal efficacy synthesis. In that setting indirectness is not a matter of degree but the central problem: the model is an acute arterial ligation in a healthy young mouse and the target is chronic ischemia in an older patient with diabetes and atherosclerosis. In place of a formal rating we report the components a reader would use to form one, namely risk of bias by domain, heterogeneity with a prediction interval, three independent small-study analyses, and a sensitivity table. An adapted framework for rating certainty in bodies of preclinical animal evidence has been described,^101^ and its domains point to the same low rating. The certainty attaching to the pooled magnitude is therefore low, mainly because of indirectness and small-study effects rather than because the direction of benefit is in doubt.

### The Pooled Magnitude, Not Its Direction, Was Sensitive to Its Underlying Assumptions

Across the sensitivity analyses (**Table 2**), treating unstated dispersions as standard error rather than standard deviation moves g from 2.26 to 1.88. Restricting to CAMARADES ≥5 gives 2.08. Restricting to studies both randomized and blindly assessed leaves 22 comparisons and an interval that no longer crosses zero (1.89, 95% CI 0.73 to 3.05). The interval is wide because the subset is small, and it should not be read as evidence of no effect. It is a fair statement of how little of this literature meets both standards at once. Leave-one-out analysis identified two studies that move the estimate by more than 0.1 (**Supplemental Figure S5**), and removing the most influential moves g from 2.26 to 1.99.

**Table 2.** Sensitivity analyses.

| Analysis | k | Hedges' g | 95% CI | Change from primary |
| --- | --- | --- | --- | --- |
| Primary three-level model with cluster-robust variance | 114 | 2.258 | 1.723 to 2.793 |  |
| Assume every unstated dispersion is SEM rather than SD | 114 | 1.883 | 1.504 to 2.263 | -0.375 |
| Exclude Liu 2013, the single most influential study | 112 | 1.994 | 1.610 to 2.379 | -0.264 |
| Treat Liu 2013 dispersions as SEM rather than SD | 114 | 2.013 | 1.635 to 2.391 | -0.245 |
| Exclude Lim 2021, whose error bars cross zero | 111 | 2.259 | 1.706 to 2.812 | +0.001 |
| Studies describing randomized allocation only | 47 | 2.720 | 1.435 to 4.004 | +0.462 |
| Studies describing both randomized allocation and blinded assessment | 22 | 1.889 | 0.728 to 3.051 | -0.369 |
| CAMARADES reporting score of 5 or more | 53 | 2.077 | 1.450 to 2.704 | -0.181 |
| Peer-reviewed full papers only, excluding conference abstracts and preprints | 101 | 2.184 | 1.617 to 2.750 | -0.074 |
| Exclude comparisons whose digitized error bars carried a high decoding risk | 104 | 2.291 | 1.706 to 2.876 | +0.033 |
| Detection-validity safeguard score of 2 or more | 82 | 2.523 | 1.672 to 3.374 | +0.265 |
| Drop the three studies from one laboratory (Cochrane 2017, Kelaini 2018, Vila-Gonzalez 2019) | 108 | 2.311 | 1.729 to 2.893 | +0.053 |
| Drop Su 2018 and Green 2024, the two largest single effects | 112 | 2.279 | 1.717 to 2.841 | +0.021 |
| Laser Doppler ratio outcomes only, excluding indocyanine green and percentage scales | 63 | 1.837 | 1.293 to 2.380 | -0.421 |
| Split the control sample size across arms sharing one control | 114 | 2.082 | 1.592 to 2.573 | -0.176 |

k is the number of comparisons. The primary model is the three-level random-effects model
with cluster-robust variance. Change is the difference in Hedges' g from the primary estimate,
computed before rounding. CAMARADES indicates Collaborative Approach to Meta-Analysis
and Review of Animal Data in Experimental Studies; CI, confidence interval; SD, standard
deviation; and SEM, standard error of the mean. Each row is refitted on the subset it names.
The randomized-allocation row retains the single comparison whose reported dispersions give
a standardized mean difference of 41.9, which is why it sits well above the primary estimate.

## Discussion

This systematic review of 68 preclinical studies finds that pluripotent stem cell-derived vascular cells improve limb perfusion in animal models of ischemia, with a pooled Hedges’ g of 2.26. The benefit was consistent in direction. Every sensitivity analysis returned a positive estimate, and the secondary outcome of limb preservation favored cell therapy. The size of the benefit is less certain. A standardized mean difference of that size places the average treated animal above roughly 99% of controls, and the 95% prediction interval, from −1.47 to 5.98, does not constrain what a new experiment would find. Egger’s regression and rank correlation indicate small-study effects, trim-and-fill moves the estimate to a little over half its unadjusted value under both of its estimators, and the funnel-plot regression extrapolates to an intercept indistinguishable from zero. That signal is not wholly one of suppression: a regression test that uses sample size in place of standard error, avoiding the mechanical correlation between a standardized mean difference and its standard error, found no significant asymmetry (P=0.80), so part of the funnel asymmetry is an artifact of the effect measure itself. Two further analyses argue that the benefit is not an artifact of the measure alone. On the natural scale the treated limb recovered about twice the perfusion of its own control (ratio of means 1.96, 95% CI 1.66– 2.33), and the pooled estimate held at fixed follow-up windows (day 14, g 1.85; day 28, g 2.19). The direction of benefit therefore survives every reasonable change of metric and timepoint, while its magnitude does not. Restricting to the 22 comparisons from the 8 studies that are both randomized and blindly assessed leaves g 1.89 (95% CI 0.73 to 3.05), an interval that excludes zero but is wide because so few studies meet both standards at once. These features are common in preclinical literatures shaped by selective reporting and unblinded measurement, and have been documented before in animal models of stroke.^6^

The mechanistic rationale on which the field is based is not yet demonstrated by its own data. Studies in which donor cells were graded as structurally incorporating into vessels reported the same benefit as studies in which they were not (P=0.30). This is the moderator with the most adjudication behind it, and a positive association would have been expected if benefit depended on donor cells becoming vessels. What the cells were shown to do is more modest than the language of the field implies: of 68 studies, 35 resolved a donor cell to a vessel wall and 32 of those walls also contained host cells, five showed a donor-only vessel, three of them exclusively, and flow through a donor-containing conduit was shown in just seven. The dominant demonstrated biology is mosaic incorporation into vessels that the host is also building, which is a real contribution to repair and may be sufficient to account for the observed benefit. Which of these patterns (donor-only vessels, mosaic incorporation, or paracrine support of host vessels) is therapeutically preferable is unknown, and the null fate-grade moderator indicates that a product need not achieve donor-only vessels to be effective.

The largest effects were seen where the mechanistic evidence was least resolvable. Studies claiming incorporation their images could not resolve reported g 3.23, against 2.33 for studies that resolved incorporation and 1.51 for those showing none. This ordering is the reverse of what the mechanistic hypothesis predicts, and it is what would be expected if the same laboratory practices that produce unresolvable images also produce unblinded, favorably read perfusion measurements. This analysis is exploratory and does not survive multiplicity adjustment. It is consistent with the small-study effects and with the null fate-grade moderator, and it sharpened rather than softened when the disputed figures were read directly. It therefore bears on how imaging evidence is generated and reported rather than on whether the therapy is effective.

The functional outcomes point the same way and are encouraging, although they are based on less evidence. Where limb status could be counted, in eleven studies, preservation strongly favored cell therapy. However, 43 of 68 studies report no functional or limb outcome at all, and behavioral testing of any kind appears in four. Limb salvage is the clinical endpoint of interest, and it is measured in barely a sixth of these studies. Perfusion therefore carries most of the evidence base, and better reporting of that surrogate would strengthen the case for translation.

A further consideration lies beyond internal validity. The target indication is chronic limb-threatening ischemia in older patients with diabetes and atherosclerosis, whereas the evidence base is overwhelmingly acute surgical ligation of a femoral artery in a healthy young mouse that recovers spontaneously, with 60 of 68 studies using healthy young animals. Effects were no smaller in the studies that used comorbid or aged animals, although only 8 studies used such a model, so the comparison has little power to detect a difference, and the acute model may not represent chronic disease. Interpretation is further constrained because most outcome values had to be digitized from figures, with only 19 of 68 studies reporting perfusion numerically. Perfusion was also measured on several non-commensurable scales, so the pooled standardized mean difference is best read as a direction and a relative magnitude rather than a literal effect size, which is one reason a modality-restricted estimate is reported. The five-level donor-cell fate grade was prespecified, whereas the expanded vascular-contribution extraction and the incorporation strata derived from it were added after registration and are reported as post hoc, with the full grading chain deposited so that any decision can be reconstructed and disputed. Because fate grade is a study-level attribute, a null association across studies constrains but does not exclude a mechanism operating within individual experiments. The analysis therefore shows that the literature does not yet provide the evidence its mechanism predicts, not that no such mechanism exists. The gap is not peculiar to limb ischemia. Across cell-based therapeutics the recurring obstacles are the choice of cell source and the demonstration that the delivered product is viable and potent where it is needed, and progress has depended on being explicit about which of these a given experiment establishes.^102^

In conclusion, pluripotent stem cell-derived vascular cells improved limb perfusion in animal models of limb ischemia, consistently in direction and with supporting evidence on limb preservation. The magnitude of that benefit remains uncertain, because the pooled estimate is large, is sensitive to reasonable changes in assumption, is unexplained by any prespecified moderator, and is accompanied by funnel asymmetry that a sample-size–based test does not reproduce, and structural incorporation of donor cells is not yet established as the mechanism. Three changes would make the next generation of studies directly informative. First, blinded assessment of the perfusion outcome, which fewer than one study in six now reports. Second, species-specific detection of donor cells in the ischemic limb, alongside a host counterstain and with demonstration of flow through a donor-containing conduit rather than mere membership of a vessel wall, and preferably in comorbid or aged animals. Third, a product definition consistent enough to be compared across laboratories and carried into trials. Adopted together, these three items would constitute a minimum reporting standard against which future preclinical studies of this therapy could be judged. With those changes, a therapy that already shows a consistent preclinical benefit can be evaluated on evidence that matches the strength of its clinical rationale.

## Acknowledgments

The authors take responsibility for the accuracy, validity, and originality of the content.

## Author Contributions

Conceptualization, M.S. and K.C.; Methodology, M.S. and K.C.; Investigation, M.S. and K.C.; Formal Analysis, M.S. and K.C.; Writing – Original Draft, M.S. and K.C.; Writing – Review & Editing, H.C., T.K., J.P.J., and S.J.; Funding Acquisition, K.C.

## Supplemental Material

-Supplemental Figures S1–S5

-Supplemental Tables S1–S5

-Completed PRISMA 2020 checklist

## Non-standard Abbreviations and Acronyms

CAMARADES: Collaborative Approach to Meta-Analysis and Review of Animal Data in Experimental Studies
ARRIVE: Animal Research: Reporting of In Vivo Experiments
CRVE: Cluster-robust variance estimation
ESC: Embryonic stem cell
GRADE: Grading of Recommendations Assessment, Development and Evaluation
iPSC: Induced pluripotent stem cell
MSC: Mesenchymal stromal cell
OSF: Open Science Framework
PAD: Peripheral artery disease
PRISMA: Preferred Reporting Items for Systematic Reviews and Meta-Analyses
PSC: Pluripotent stem cell
SYRCLE: Systematic Review Centre for Laboratory Animal Experimentation

